# Identification and Characterization of PCAHIT, a Novel Long Non-Coding RNA Associated with Poor Outcomes in Prostate Cancer

**DOI:** 10.1101/2025.05.21.655369

**Authors:** Hui Li, Noah S. Younger, Bhavna Malik, Hyun Jin Shin, Chao Zhang, Yashar Niknafs, Shuang (George) Zhao, Kari Wilder-Romans, Sethuramasundaraman Pitchiaya, Jasmine Anderson, Sumin Han, Travis Barnard, Paul Lloyd, Meng Zhang, Lisa N. Chesner, Marsha Calvert, Emily A. Egusa, Jun Zhu, Jonathan Chou, Rajdeep Das, Vishal Kothari, Tanu Shenoy, Morgan E. Diolaiti, Rohit Malik, John R. Prensner, Alma Burlingame, Alan Ashworth, Arul M. Chinnaiyan, Felix Feng, Haolong Li

## Abstract

**Background:** Despite extensive investigation, the factors promoting aggressive prostate cancer are poorly understood. In particular, despite a few prominent examples, the role of long non-coding RNAs (lncRNAs) is largely unknown.

**Methods:** We performed a comprehensive analysis of whole-genome transcriptome data to identify differential expression across 1,567 patients with prostate cancer, then correlated differential expression with risk of metastatic recurrence. We characterized the lncRNA most associated with metastasis *in vitro* and *in vivo* to investigate the mechanism by which it promotes aggressive prostate cancer.

**Results:** We have identified a novel lncRNA, Prostate Cancer Associated hnRNPK Interacting Transcript (PCAHIT), which is strongly associated with metastasis and poor overall survival in men with prostate cancer. We find that overexpression/knockdown of PCAHIT in pre-clinical models of prostate cancer modulates proliferation rates and markers of an aggressive phenotype through regulation of steroid biosynthesis and expression of the MHC class I complex, driving increased growth in androgen-depleted conditions and decreased susceptibility to T cell-mediated cytotoxicity. We further find that the heterogeneous nuclear ribonucleoprotein hnRNPK directly binds PCAHIT and is indispensable for its activity.

**Conclusions:** Overall, our findings indicate that PCAHIT is a driver of aggressive prostate cancer phenotypes and poor clinical outcomes through suppression of the immune response and increased androgen-independent cancer growth. PCAHIT is a potential biomarker of aggressive disease.

## Introduction

Prostate cancer is the most common non-cutaneous malignancy in men, accounting annually for approximately 268,000 cases and 35,000 deaths in the United States.(1) The vast majority of patients are diagnosed at an early, localized stage and undergo either prostatectomy or radiation therapy with curative intent. Of these patients, a subset will recur and go on to develop metastatic disease, which is incurable. Historically, clinical characteristics such as pre-treatment PSA level, disease extent, and tumor Gleason score have been used to predict risk of recurrence,(2) however, the poor predictive ability of these biomarkers has motivated a search for genetic factors that may augment or replace existing models. These efforts have achieved notable success in identifying multiple protein coding genes which are drivers of aggressive disease including *PTEN,*(3) *MYC,*(4) *EZH2*(5) and others.(6),(7),(8) These findings have begun to influence clinical practice with development of the Decipher, Oncotype DX, and Prolaris scores, all of which are multi-gene assays to predict recurrence risk after localized therapy and have recently been incorporated into national guidelines for management of prostate cancer.(9) The identification and validation of molecular drivers of prostate cancer progression has also contributed to our understanding of prostate cancer disease biology. Despite these successes, our knowledge of factors driving aggressive biology and risk of recurrence remains incomplete, and there is great need for the discovery of novel predictive biomarkers of aggressive disease to determine which patients might benefit from additional treatments and closer monitoring.

Over 70% of the human genome is composed of non-coding elements, the function of which is largely unknown.(10) Long non-coding RNAs (lncRNAs) are RNA transcripts of over 200 nucleotides that are not transcribed into proteins. lncRNAs are widely expressed, and by virtue of base pair interactions and unique tertiary structure can form specific RNA-RNA, RNA-DNA, and RNA-protein interactions that are important for biology and disease. Although most studies investigating drivers of prostate oncogenesis have focused on protein coding genes, it has become increasingly clear that lncRNAs play a critical role in gene regulation in both normal and cancerous cells. Examples from prostate cancer include ARLNC1,(11) which increases AR signaling by stabilizing the Androgen Receptor (AR) gene transcript, CTBP1-AS,(12) which antagonizes expression of multiple tumor suppressor genes through recruitment of the transcriptional repressor PSF to target loci, PCAT-1,(13),(14) which promotes prostate cancer proliferation by repressing transcription of tumor suppressor genes including *BRCA2*, PCAT- 19,(15) which activates a subset of cell-cycle genes to promote prostate cancer progression, and SChLAP1,(16) which antagonizes the chromatin-modifying SWI/SNF complex and promotes invasion and metastasis. Despite a few well-characterized examples, the vast majority of the estimated 58,648 lncRNAs have only been annotated in the past decade and remain poorly understood. In order to investigate the role of lncRNAs in prostate cancer progression and metastasis, we analyzed microarray data from a large cohort of patients with localized prostate cancer and long-term follow-up to identify differential expression of lncRNAs associated with increased risk of metastatic recurrence. This analysis nominated the novel lncRNA we name PCAHIT (Prostate Cancer Associated hnRNPK Interacting Transcript) as being most significantly associated with recurrence risk. We investigated the function of this lncRNA to clarify the mechanisms by which it promotes aggressive prostate cancer phenotypes.

## Materials and Methods

### Microarray

Microarray data was obtained from previously published datasets(17–22) to create a pooled cohort of 1,567 samples. Microarray processing and normalization was performed as described previously.(23, 24) Gene expression of all annotated and un-annotated lncRNAs(25) was calculated by taking the mean expression of probesets mapping to exons. Single exon lncRNAs, or genes with a 95^th^ percentile RNA-seq expression in prostate < 1 FPKM(25) were excluded. Nomination was performed by putting each gene into a Cox model with expression as a continuous variable in the pooled cohort. The lncRNAs were then ranked by the C-index. Mapping of probesets to PCAHIT was then refined by using the sequence from RACE.

### Cell lines

LNCaP, C4-2B, PC-3, VCaP, 22Rv1, MDA-PCa-2b, RWPE and DU-145 cells were purchased from ATCC, grown in respective ATCC recommended culture media, and maintained at 37°C in 5% CO2. All cell lines were regularly tested for Mycoplasma contamination using MycoAlert Mycoplasma Detection Kit (Lonza #LT07-318). For androgen-deprivation studies, LNCaP cells were grown in phenol red free RPMI media supplemented with 5% charcoal-stripped serum (Thermo Fisher Scientific #12676029). LNCaP-AR-enza-res cells were generated by treating LNCaP-AR parental cells with increasing doses of enzalutamide (from 1 mmol/L to 50 mmol/L) until resistance emerged.

### siRNA, ASO and shRNA mediated knockdown

Prostate cancer cell lines were transfected with siRNA (Dharmacon), ASO (Qiagen) or shRNA (this study) targeting PCAHIT, hnRNPK, or control non-targeting using Lipofectamine RNAiMAX (Invitrogen #13778100) at a final concentration of 50 nM according to the manufacturer’s instructions. Cells were collected and used for downstream experiments 48 hours following transfection. Sequences are as follows:

- siRNA PCAHIT-1: 5’ - CCAUAUACCUGAUGCAGAAUU – 3’
- siRNA PCAHIT-2: 5’ - AAAUAAACUCCAAGCAGAAUU – 3’
- shRNA PCAHIT-1: 5’ - AGAAGAGAATGTACA – 3’
- shRNA PCAHIT-2: 5’ - AAATAAACTCCAAGA – 3’
- ASO-PCAHIT-1: 5’ - GTTAGTCACATGGATA – 3’
- ASO-PCAHIT-2: 5’ - AAAGGTGTGCTCGGA – 3’
- siNTC (Dharmacon Non-targeting Control Catalog: D-001810-10-20)
- sihnRNPK-1 (Dharmacon HNRNPK siRNA Catalog: J-011692-05-0002)
- sihnRNPK-2 – (Dharmacon HNRNPK siRNA Catalog: J-011692-06-0002)

### Rapid amplification of cDNA ends (RACE)

RACE was performed as previously described.(26) Briefly, 5’ and 3’ RACE was performed using the GeneRacer RLM-RACE kit (Invitrogen #L1500-01) according to the manufacturer’s protocols and using Platinum Taq high-fidelity polymerase (Invitrogen #15966005) for PCR.

Following resolution on a 1.5% agarose gel, individual bands were excised and cloned into PCR4 TOPO vector (ThermoFisher #450030), then sequenced.

### *In vitro* RNA-binding assay and mass spectrometry

The RNA-mediated protein pull down assay was performed using the RiboTrap Kit (MBL International #RN1001) standard protocol. Briefly, 5-bromo-UTP (BrU) was randomly incorporated into sense PCAHIT, antisense PCAHIT control, and LacZ mRNA control via T7 *in vitro* transcription. Then the BrU incorporated RNA samples were bound to beads conjugated with anti-BrdU antibodies and incubated with nuclear fractions isolated from LNCaP or C42B cells. The beads were washed to remove non-specific interactions, bound proteins were eluted and collected, then alkylated by the addition of iodacetamide at a concentration of 15 mM. The eluted samples were subjected to SDS–PAGE electrophoresis and individual lanes were excised and digested with trypsin for 2 hours. Samples were then sent to the UCSF proteomic core facility for mass spectrometry, which was performed using a Velos Pro mass spectrometer (Thermo Scientific) operated in data-dependent mode using 120-min gradients in EASY-LC system (Proxeon) with 95% water, 5% acetonitrile (ACN), 0.1% formic acid (FA) (solvent A), and 95% ACN, 5% water, 0.1% FA (solvent B) at a flow rate of 220 nl min^−1^.

### Invasion assay

Invasion assays were performed as previously described.(26) Briefly, 3 x 10^5^ cells were seeded in a 24-well FluoroBlok chamber (Corning #351152) pre-coated with Matrigel (Corning #354237) in culture media supplemented with charcoal-stripped serum. Culture media with 10% FBS was used in the lower chamber. After 48 hours, cells in the lower chamber were stained with calcein AM (Invitrogen #C34852) and quantified using a fluorescent microscope.

### Quantitative RT-PCR

Total RNA was isolated from cells using the Quick RNA Kit (Zymo Research #R1054); cDNA was synthesized using the Superscript III RT First Strand Kit (Invitrogen #18080051). Quantitative polymerase chain reaction was performed using Fast SYBR Green master mix (Thermo Fisher Scientific #4385612) on an Applied Biosystems’ QuantStudio7. Ct values were normalized to GAPDH, and relative expression was calculated using the 2^ddCt^ method. All experiments were performed at least in triplicate. Primer sequences are from the Harvard PrimerBank:

- PCAHIT

- Fwd: 5′ - aaagatgtgagagaattcaactgat – 3′
- Rev: 5′ - tggcatgacagaggaaacaa – 3′
- SChLAP1

- fwd: 5′ - tggacacaatttcaagtcctca – 3′
- Rev: 5′ - catggtgaaagtgccttataca – 3′
- AKR1C3

- fwd: 5′ - ccgaagcaagattgcagatggc- 3′
- rev: 5′ - gtgagttttccaaggctggtcg– 3′
- UGT2B10

- Fwd: 5′ - ccagatgccttaggtctcaatac– 3′
- Rev: 5′ - gcctcatagatgccattggctc– 3′
- UGT2B15

- Fwd: 5′ - actttaggttccaatactcgactg– 3′
- Rev: 5′ - cgcctcatagatgccattggttc– 3′
- TMPRSS4

- Fwd: 5′ - gacgaggagcactgtgtcaaga– 3′
- Rev: 5′ - gaaacaggcagagaaccagttcc– 3′
- XIST

- Fwd: 5′ - gaaccacctacacttgag– 3′
- Rev: 5′ - tgctatgcgttatctgag– 3′
- HLA-A

- Fwd: 5′ - gaccaggagacacggaatgt– 3′
- Rev: 5′ - gatgtaatccttgccgtcgt– 3′
- HLA-B

- Fwd: 5′ - gtattgggaccggaacacac– 3′
- Rev: 5′ - gatgtaatccttgccgtcgt– 3′
- HLA-C

- Fwd: 5′ - atacctggagaacgggaagg– 3′
- Rev: 5′ - gaaggttccatctcctgctg– 3′
- U1

- Fwd: 5′-caggggagataccatgatcacgaag-3′
- Rev: 5′-ggtcagcacatccggagtgcaatgg-3′

### RNA Sequencing

22Rv1 cells were transfected with pCDH plasmid (Addgene #72265) for overexpression of PCAHIT or empty vector control. RNA was extracted from the cells as described above and QuantSeq 3′mRNA-Seq library prep kit FWD for Illumina (Lexogen, Cat# 015.24) was used to prepare the library as per the manufacturer’s protocol. Quality control was performed by using the Agilent Bioanalyzer 2100 system and the samples were sequenced using an Illumina HiSeq 4000. Two independent biological replicates were performed for each condition.

### GSEA

The gene set enrichment analysis was performed using the pre-ranked method implemented in the fgsea R package (DOI: 10.18129/B9.bioc.fgsea),(27) and KEGG gene sets were downloaded from the molecular signatures database (MSigDB); genes were ranked by the Wald-statistics from DESeq2. Normalized enrichment scores quantified over-/under- representation of predefined gene sets in cells overexpressing PCAHIT versus control.

### Immunoblotting

Cells were lysed in RIPA buffer containing Halt protease and phosphatase inhibitor cocktail (Thermo Scientific). Lysates were subjected to SDS-PAGE gel electrophoresis, transferred to poly(vinylidene fluoride) membranes, blocked in 5% w/v bovine serum albumin, and incubated with primary antibody overnight at 4°C. The blot was visualized using ECL Detection Reagents (Genesee Scientific). Antibodies against vinculin (Cell Signaling Technology #13901), AKR1C3 (Sigma-Aldrich A6229), UGT2B10 (Novus Bio NBP2-93266), UGT2B15 (Novus Bio NBP2- 94747), MHC I (Origene, AM33035PU-N), hnRNPK (Cell Signaling Technology #4675), PTBP1 (MBL Life Science #RN011P), and nucleolin (Cell Signaling Technology #14574) were used according to the manufacturer’s recommended dilutions.

### Nuclear/Cytosolic lysates

Cellular fractionation was performed as previously described(26) using a NE-PER Nuclear and Cytoplasmic Extraction Kit (Thermo Fisher #78833), according to the manufacturer’s instructions. RNA was isolated and qRT–PCR was performed as described above.

### Cell growth and scratch-wound assay

For the growth assay, cells were seeded in a 96-well cell culture plate in growth medium at a density of 2 × 10³ cells/well. Cell growth, measured by confluence or indicated by the Nuclight Red signal in each well, was monitored daily for 7 days. Images were acquired and analyzed using the IncuCyte® S3 Software (Sartorius).

For the scratch wound assay, cells were seeded at a high density of 25,000 cells/well in 96-well plates. Once confluent, the cells were scratched using the IncuCyte 96-Well Wound Maker tool (Sartorius) and washed once to remove floating cells and prevent reattachment in the wound area. Images were captured and analyzed using the IncuCyte S3 Live-Cell Analysis System (Sartorius).

### WST-1 Assay

22Rv1 control and 22RV1 PCAHIT cells were seeded in a 96-well plate in CSS and 5mM enzalutamide. 3 days and 7 days after seeding, cells were incubated with Cell Proliferation Reagent WST-1 (Roche #5015944001). Absorbance values were obtained using an Epoch Microplate Spectrophotometer and analyzed using Prism8 (GraphPad).

### Mouse xenograft

*In vivo* experiments were performed as previously described.(26) A total of 5 × 10^6^ cells of C42B control or C42B shPCAHIT cells suspended in 100 μl of PBS/Matrigel (Corning #356230) (1:1) were injected subcutaneously in 5-week-old pathogen-free male CB-17 severe combine immunodeficient mice (CB-17 SCID). Tumors were measured weekly using a digital caliper and tumor volumes were estimated using the formula (π/6) (*L* × *W*2), where *L*=length of tumor and *W*=width. Mouse lungs were collected to determine spontaneous metastasis by measuring human Alu sequence. Briefly, genomic DNA from lungs were prepared using Puregene DNA purification system (Qiagen #158667), followed by quantification of human Alu sequence by human Alu specific Fluorogenic Taqman qPCR probes.

### Single-molecule fluorescence *in situ* hybridization

FISH was performed as previously described.(26) Briefly, LNCaP cells were grown on 8-well chambered coverglass, formaldehyde fixed and permeabilized overnight at 4°C using 70% ethanol, rehydrated in 10% formamide and 2x SSC for 5 minutes, then treated with 10 mM pooled FISH probes for 16 hours in 2x SSC containing 10% dextran sulfate, 2 mM vanadyl-ribonucleoside complex, 0.02% RNAse-free BSA, 1 µg µl^-1^ *E. coli* transfer RNA and 10% formamide at 37°C. Cells were then washed twice for 30 minutes at 37°C with 10% formamide in 2x SSC and mounted in 10 mM Tris/HCl pH 7.5, 2x SSC, 2 mM trolox, 50 µM protocatechiuc acid and 50 nM protocatechuate dehydrogenase. Samples were imaged using HILO illumination. Probe sequences are listed in **Table S1**.

### T Cell cytotoxicity assay

The T cell cytotoxicity assay was performed as previously described. Briefly, 22Rv1 cells with endogenous expression of the cancer-testis antigen NY-ESO1 and transduced with Nuclight lentivirus reagent were seeded at 2,500 cells per well in 96-well plates (Falcon #353072) and allowed to attach overnight. T cells expressing an HLA-A2-restricted TCR specific for NY-ESO1 were then added at a 1:1 E:T ratio. RFP-expressing target cells were imaged and counted every 6 hours for 3 days using an Incucyte.

### Statistics

For analysis of microarray data, high/low gene expression was determined by splitting on the median expression level. Kaplan-Meier curves are shown and statistical inference was performed using the Log-rank test. All statistical tests were two-sided and significance was set as p<0.05. All analyses were performed in R 3.1.2. Other statistical analysis was performed using GraphPad Prism 9.0 software. Data are presented as mean SEM, and mean differences between groups were compared using two-way ANOVA. N is ≥ 3 independent replicates for each experiment.

## Results

### Nomination of PCAHIT as a lncRNA Associated with Poor Prognosis in PCa

We previously analyzed 7,256 RNA sequencing libraries from tumors, normal tissues, and cell lines from 25 independent studies, and identified 58,648 lncRNAs, of which 48,952 (79%) are novel and previously unreported.(25) We assessed the prognostic significance of these lncRNAs using high-density microarray data from a cohort of 1,567 patients with a history of localized PCa treated with prostatectomy who then underwent long-term follow-up. This analysis nominated a novel lncRNA we name PCAHIT (Prostate Locus of Uncharacterized Transcript Outlier 201) as more strongly associated with metastatic progression in this cohort than any other protein-coding or non-coding gene (**Figure 1A, B**). High PCAHIT expression is significantly associated with poor metastasis-free survival (hazard ratio 1.7, p-value <1x10^-7^) (**Figure 1C**) and overall survival (hazard ratio 1.4, p-value <0.001) (**Figure 1D**). Notably, average expression of PCAHIT in prostate cancer samples is higher than that of the recently characterized lncRNA PCAT-19 (average RPKM of 0.27 in mCPRC).(28)

**Figure 1:**
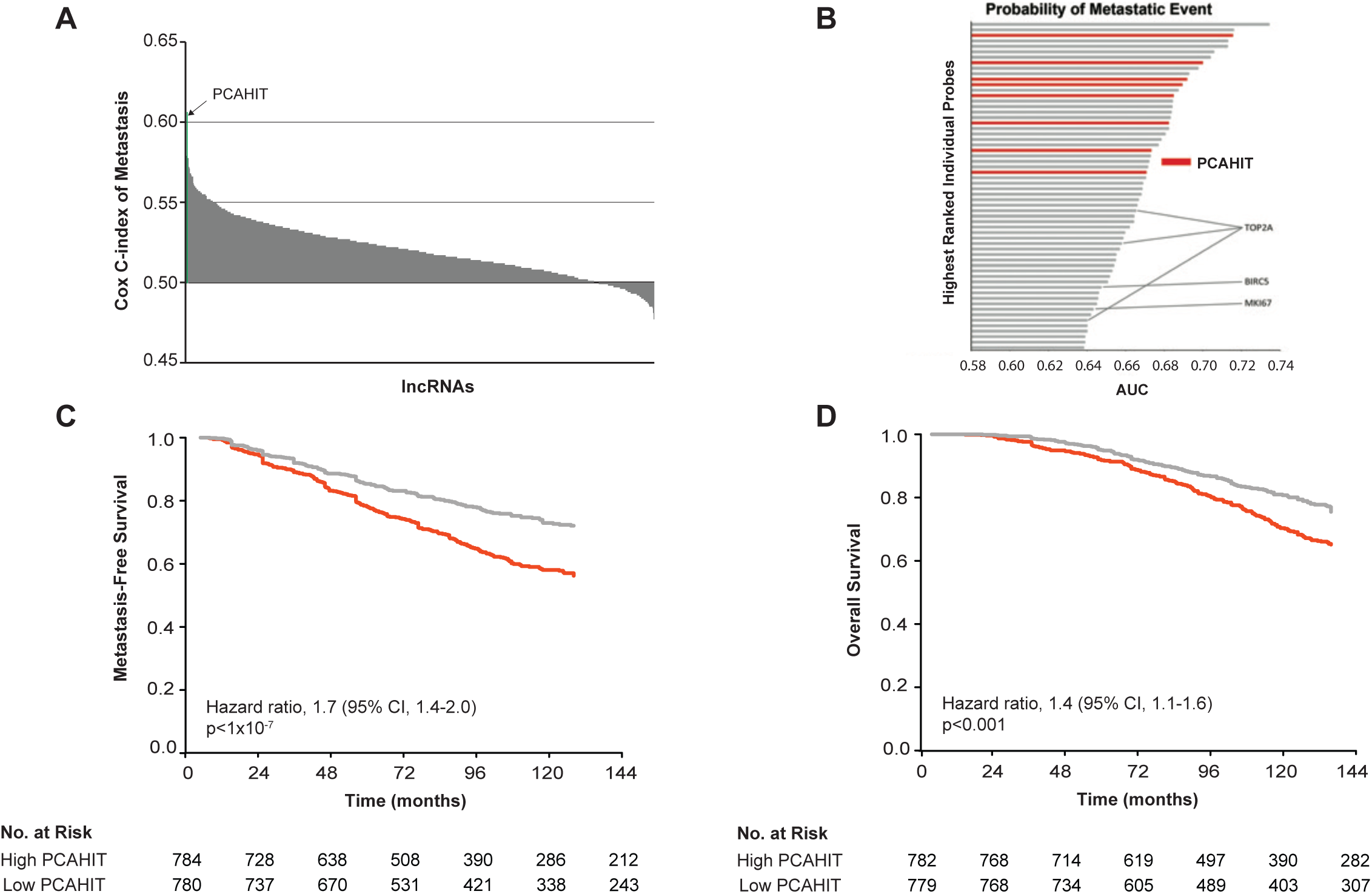
The novel lncRNA PCAHIT is associated with prostate cancer progression, metastasis and poor prognosis. Analysis of microarray data with associated clinical outcomes data from a cohort of 1,567 patients with localized prostate cancer undergoing prostatectomy. (A) Rank ordering of all lncRNAs by Cox-C index of association with metastasis, from 1567 patient cohort with initial diagnosis of localized PCa treated with prostatectomy and subject to high-density microarray analysis. (B) The probability of metastasis events calculated for each individual probe from the array (including protein coding genes) and ranked from high to low. Probes against PCAHIT are highlighted in red, while those against other genes known to be associated with increased risk of metastasis (TOP2A, BIRC5, and MKI67) are marked. (C and D) Kaplan Meier plots showing metastasis-free survival (C) and overall survival (D) stratified by high- versus low-PCAHIT expression levels.

### Characterization of PCAHIT

Having identified PCAHIT as a novel lncRNA associated with poor prognosis in prostate cancer, we next sought to characterize this transcript. We first interrogated RNAseq data from the TCGA(29) and Stand Up To Cancer (SU2C)(30, 31) datasets to determine expression patterns across benign and malignant samples from multiple tissue types, finding high levels of PCAHIT expression specifically in prostate cancer (**Figure 2A**). We further characterized the expression profile of PCAHIT in a variety of prostate cancer cell lines by quantitative PCR (qPCR), finding elevated expression in LNCaP and MDA-PCa-2B cell lines compared to VCaP, PC3, DU-145, 22Rv1 or RWPE cell lines (**Supplemental Figure 1**).

**Figure 2:**
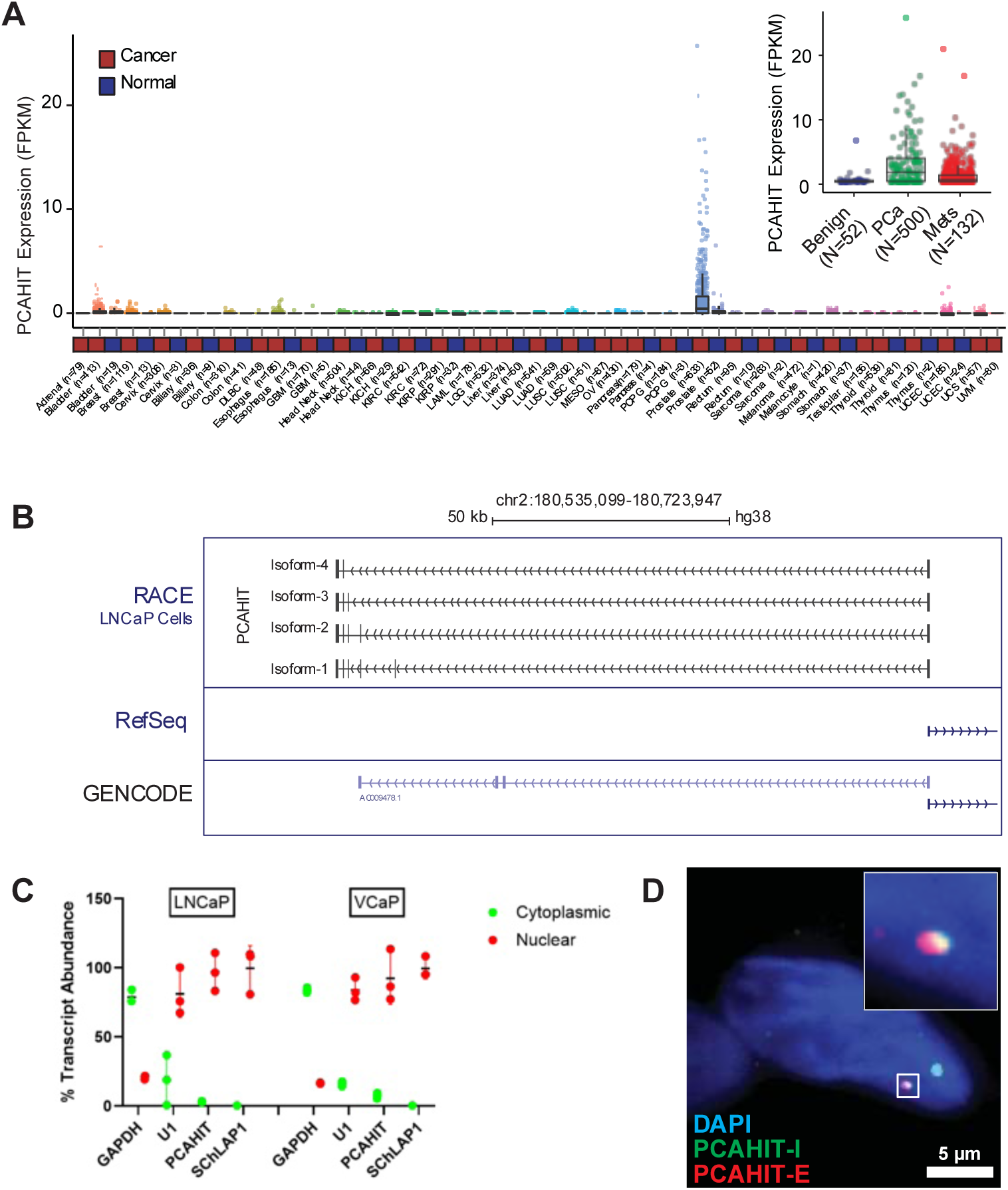
PCAHIT is a prostate cancer specific lncRNA, primarily localizing to the nucleus. (A) Expression levels (FPKM) of PCAHIT in normal tissues and localized cancers, from TCGA and SU2C datasets. (B) RACE (Rapid Amplification of cDNA Ends) assay identifying four distinct PCAHIT isoforms aligned with representation of PCAHIT in genome browser and current annotations in Refseq and GENCODE databases. (C) Quantitative rtPCR analysis of RNA isolated from the cytosolic or nuclear fraction of cultured LNCaP and VCaP prostate cancer cells (n=3). (D) Representative pseudocolored image of a LNCaP cell probed by single-molecule fluorescence in situ hybridization (smFISH) for PCAHIT introns (green) and PCAHIT exons (red). The nucleus is stained with DAPI (blue). Inset, zoomed in view of grey box, 3.3 x 3.3 μm^2^.

Analysis using the RefSeq and GENCODE databases shows that PCAHIT is a transcribed gene with no substantial (> 66 aa) open reading frame. To further delineate the gene structure of PCAHIT, we performed 5′ and 3′ RACE (Rapid Amplification of cDNA Ends) in LNCaP cells and in human samples (WA28-23 tissue) and found that PCAHIT is a 1.1kb polyadenylated transcript composed of 6 exons that are alternatively spliced to produce four major isoforms (**Figure 2B**). PCAHIT is located on the anti-sense strand of human chromosome 2q31.3 near SChLAP1, a lncRNA also associated with poor outcomes in prostate cancer.(16)

To investigate the subcellular localization of PCAHIT, we performed qPCR assays after cellular fractionation, finding that the majority (∼90%) of PCAHIT lncRNA is located in the nuclear fraction in both LNCaP and VCaP cell lines (**Figure 2C**). Single-molecule RNA FISH (smFISH) of both immature (pre-splicing) and mature (post-splicing) PCAHIT confirms the nuclear localization of PCAHIT transcripts (**Figure 2D**).

### PCAHIT Promotes Aggressive Prostate Cancer Phenotypes *In Vitro* and *In Vivo*

Having characterized PCAHIT as a lncRNA with nuclear localization, we next sought to investigate the functional importance of PCAHIT in prostate cancer. We first performed shRNA mediated knockdown of PCAHIT in LNCaP and C42B PCa cell line models, targeting exon 6 specifically, which is common to all four PCAHIT isoforms. We obtained knockdown efficiency of approximately 60% and 80% in the two cell lines, respectively (**Supplemental Figure 1**). Notably, despite the close spatial association of PCAHIT and SChLAP1, we achieve highly selective knockdown of PCAHIT compared to SChLAP1 (**Supplemental Figure 2**). In both cell lines, knockdown of PCAHIT results in a decreased proliferation rate *in vitro* (**Figure 3A,B**). To validate this finding *in vivo*, we performed subcutaneous injections of C42B cells stably transduced with shRNA targeting PCAHIT or a non-targeting control into CB17 SCID mice. Tumor growth (**Figure 3C**) and metastasis (**Figure 3D**) were significantly reduced upon PCAHIT knockdown. We next investigated the role of PCAHIT in promoting prostate cancer invasion *in vitro* using the matrigel Boyden chamber assay. C42B cells with PCAHIT knockdown show reduced invasion through the membrane relative to control (**Figure 3E**). Conversely, 22Rv1 cells overexpressing PCAHIT show an increased rate of migration by the scratch-wound assay (**Figure 3F**), evidenced by a more rapid decrease in the cell-free ‘wound’ width over time. Overall, these data suggest that PCAHIT promotes prostate cancer proliferation, invasion, and metastasis.

**Figure 3:**
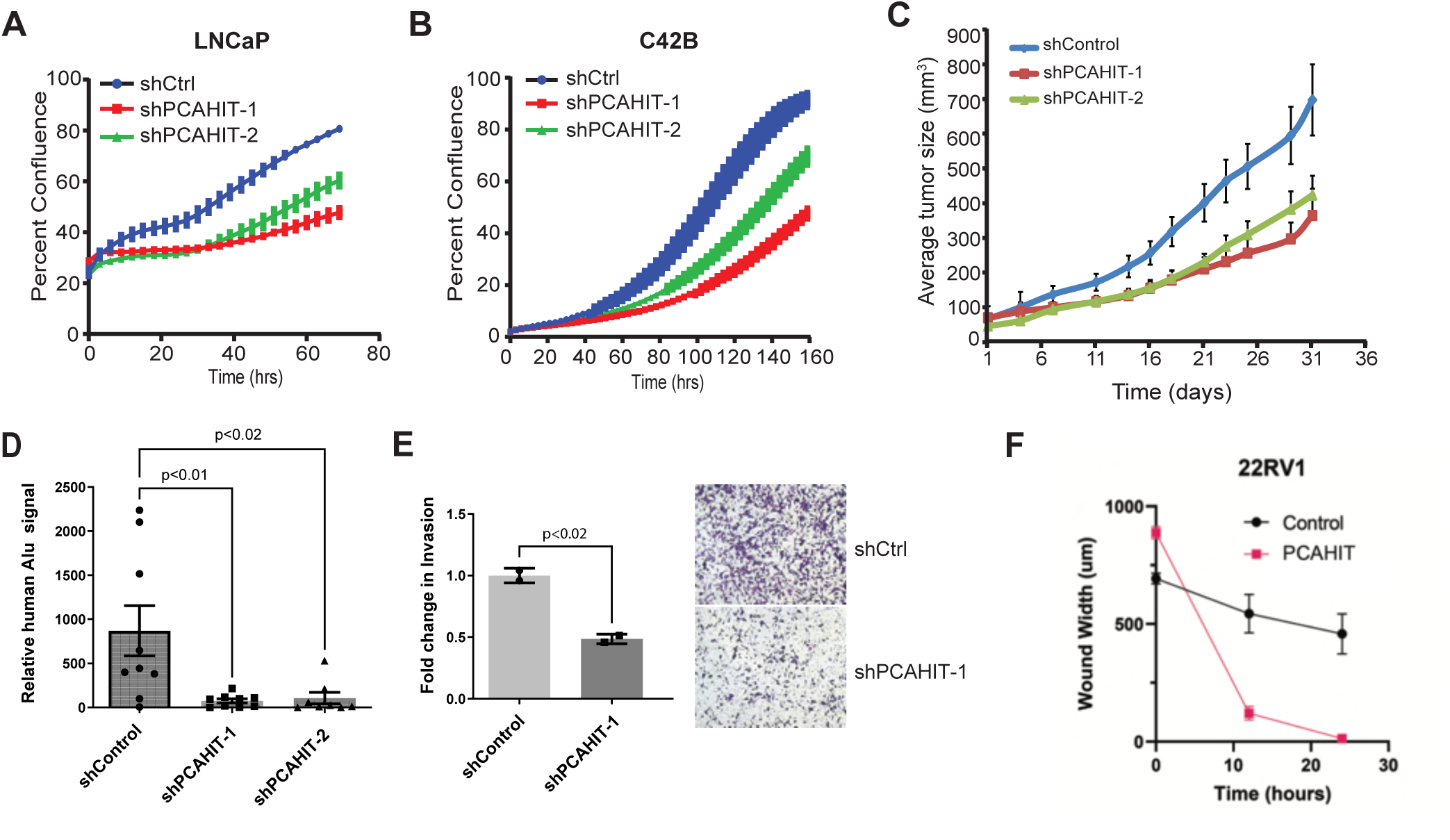
Modulating PCAHIT expression level alters aggressive phenotypes in prostate cancer models. *In vitro* growth assay in LNCaP (A) and C42B (B) prostate cancer cell lines, with short hairpin RNA (shRNA) mediated knockdown of PCAHIT (n=3). (C) *In vivo* growth of C42B cells transduced with either control or PCAHIT targeting shRNA injected subcutaneously into CB-17 SCID mice. (D) Quantification of metastases in CB-17 SCID mice by quantitative PCR amplification of human Alu signal in dissected lung tissue following subcutaneous injection of C42B cells stably transduced with either control or PCAHIT targeting shRNA (n=9, reported as mean +/- SEM). (E) Boyden chamber assay with C42B cells stably transduced with either control or PCAHIT targeting shRNA (n=2 per guide). (F) Quantification of cell migration (control or overexpressing PCAHIT) following mechanical clearance from a portion of a coverslip (n=3).

### PCAHIT Regulates MHC Class I and Steroid Biosynthesis Gene Expression

Having established a role for PCAHIT in promoting aggressive prostate cancer phenotypes, we next sought to investigate the mechanism of PCAHIT activity. Based on the nuclear localization of PCAHIT and established roles for multiple lncRNAs in transcriptional regulation,(32) we hypothesized that PCAHIT may function in the regulation of transcription in prostate cancer. We therefore performed RNA sequencing (RNA-seq), comparing wild type 22Rv1 cells to 22Rv1 cells overexpressing PCAHIT (**Figure 4A**). Gene Set Enrichment Analysis (GSEA) shows, among other pathways affected, upregulation in steroid hormone biosynthesis genes and downregulation in genes involved in multiple pathways related to immunity and inflammation (**Figure 4B**). We decided to further investigate these particular pathways because of prior work suggesting their importance in prostate cancer, specifically evasion of immune surveillance by downregulation of MHC Class I genes,(33) and resistance to therapy targeting androgen receptor signaling via cell autonomous androgen production.(34),(35),(36) Representative steroid hormone biosynthesis genes (AKR1C3, UGT2B10, UGT2B15) and MHC Class I genes (HLA-A, HLA-B, HLA-C) were selected for individual validation and qPCR shows the expected up- or down-regulation, respectively, upon exogenous overexpression of PCAHIT (**Figure 4C**). We confirmed that these changes in expression levels correlate with changes in protein levels by western blot analysis (**Figure 4D**).

**Figure 4:**
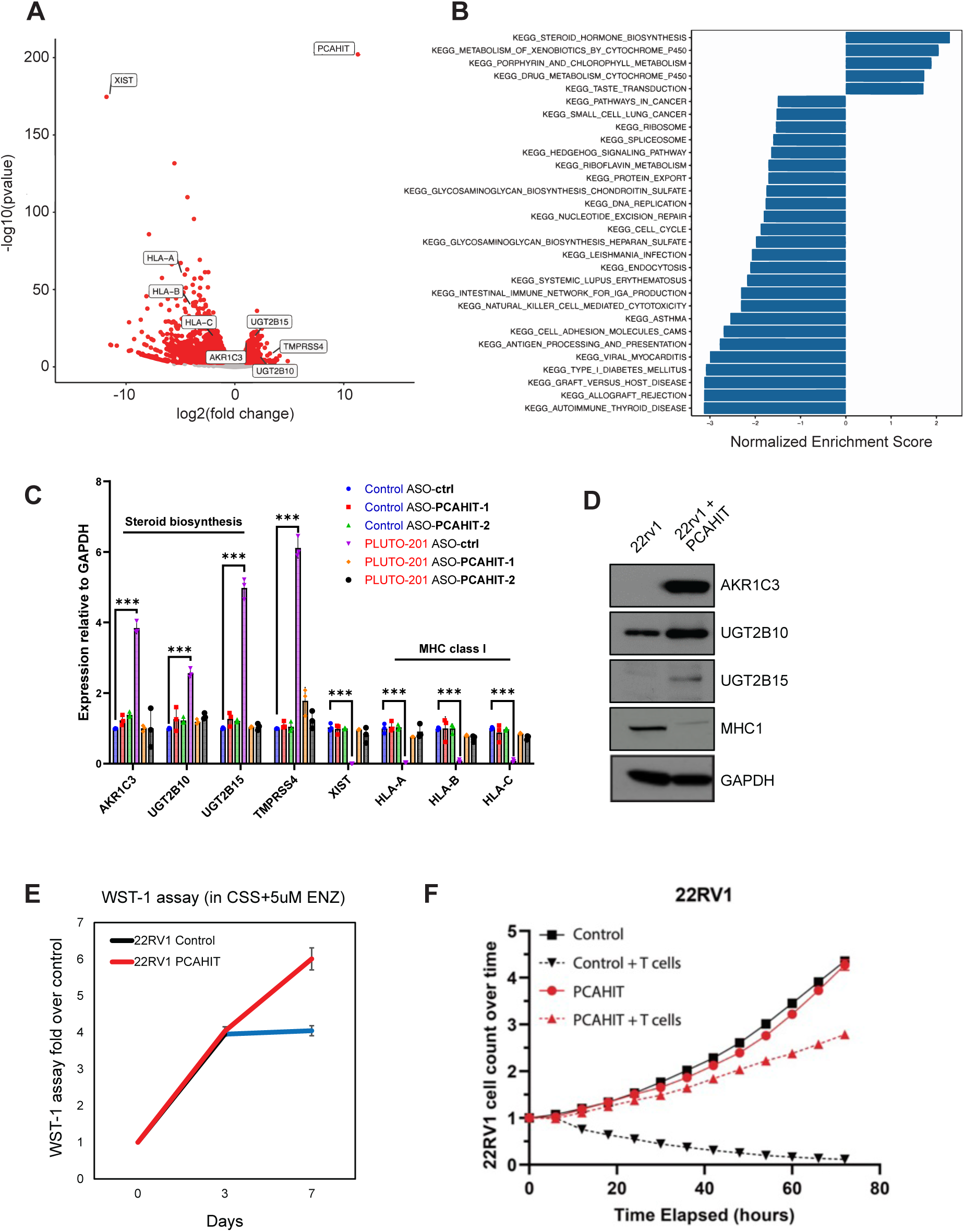
PCAHIT regulates expression of immune related genes. (A) RNA-seq from 22RV1 wt versus 22Rv1 prostate cancer cells overexpressing PCAHIT (n=2). Differentially expressed genes between the two groups are highlighted in red. (B) Gene Set Enrichment Analysis (GSEA) from RNA-seq data. (C) qPCR of putative PCAHIT regulated genes following knockdown and/or overexpression of PCAHIT (n=3). (D) Western bot of 22Rv1 wt or PCAHIT overexpressing cells for putative PCAHIT regulated genes (n=2, representative image shown). (E) Proliferation assay of 22Rv1 wt or PCAHIT overexpressing cells in androgen depleted conditions (n=3). (F) Proliferation assay of 22Rv1 wt or PCAHIT overexpressing cells in the presence or absence of T cells (n=3).

Because prostate cancer is typically dependent on androgen signaling for growth and survival,(37),(38) we hypothesized that increased PCAHIT expression may upregulate steroid hormone biosynthesis genes and increases intracellular androgen production, thereby promoting prostate cell growth in androgen depleted conditions. We tested this hypothesis by measuring proliferation rates of 22Rv1 cells overexpressing PCAHIT versus control in charcoal stripped serum supplemented with the direct androgen receptor antagonist enzalutamide, which binds competitively with androgen and inhibits androgen receptor activation.(39) We find that under these conditions, PCAHIT overexpression increases proliferation (**Figure 4E**).

Because MHC Class I genes are critical for T cell-mediated cytotoxicity,(40) we hypothesized that increased PCAHIT expression may decrease T cell-mediated cytotoxicity by downregulating MHC Class I genes. We tested this hypothesis using an *in vitro* T cell cytotoxicity assay, in which Red Fluorescent Protein (RFP) and *NY-ESO1*-expressing 22Rv1 cells are co-cultured with CD8 T cells transduced with a *NY-ESO1* T cell receptor while assessing change in 22Rv1 growth and survival by quantifying the number of surviving RFP-expressing cells. This assay confirms decreased T cell-mediated cytotoxicity upon overexpression of PCAHIT (**Figure 4F**).

Overall, these data show that PCAHIT promotes expression of steroid biosynthesis genes and represses MHC Class I genes. Consistent with the expected functional outcome of this pattern of gene regulation, PCAHIT overexpression promotes increased prostate cancer cell proliferation in an androgen depleted environment, as well as increased resistance to T cell-mediated cytotoxicity.

### hnRNPK is Required For PCAHIT Mediated Downstream Gene Regulation

After defining the transcriptional pathways important for the aggressive prostate cancer phenotype mediated by PCAHIT, we next sought to further define the mechanism of action of PCAHIT. Because lncRNAs typically regulate transcription via critical interactions with proteins such as transcription factors, chromatin modifying enzymes, and/or other RNA binding factors,(41) we attempted to identify protein binding partners using immunoaffinity purification of labeled PCAHIT followed by LC-MS/MS (RiboTrap Assay). Using the complementary PCAHIT anti-sense RNA strand as a control, we found specific interactions between PCAHIT and both heterogeneous nuclear ribonucleoprotein K (hnRNPK) and polypyrimidine tract binding protein 1 (PTBP1) (**Figure 5A-C**). Analysis of clinical data from our initial patient cohort shows that elevated hnRNPK expression in prostatectomy samples, but not elevated PTBP1 expression, is associated with worse progression-free survival (**Supplemental Figure S3A**). HnRNPK is a DNA- and RNA- binding protein involved in mRNA maturation, stability, and translational control.(42) Furthermore, hnRNPK is known to bind and regulate lncRNA function(43),(44) and is implicated in prostate cancer pathogenesis.(45) We therefore decided to focus further investigation on the role of hnRNPK in facilitating PCAHIT-mediated transcriptional changes in prostate cancer. To establish evidence of direct and specific binding between hnRNPK and PCAHIT we sought to define the region of PCAHIT which interacts with hnRNPK. Performing affinity purification with truncated PCAHIT constructs, we found that PCAHIT exon six alone is sufficient for binding of hnRNPK, while exons 1-5 show no significant interaction with hnRNPK (**Supplemental Figure S3B,C**). Existing reports suggest that hnRNPK binds preferentially to G:C-rich nucleotide sequences,(46) which in PCAHIT are predominantly found in exon six.

**Figure 5:**
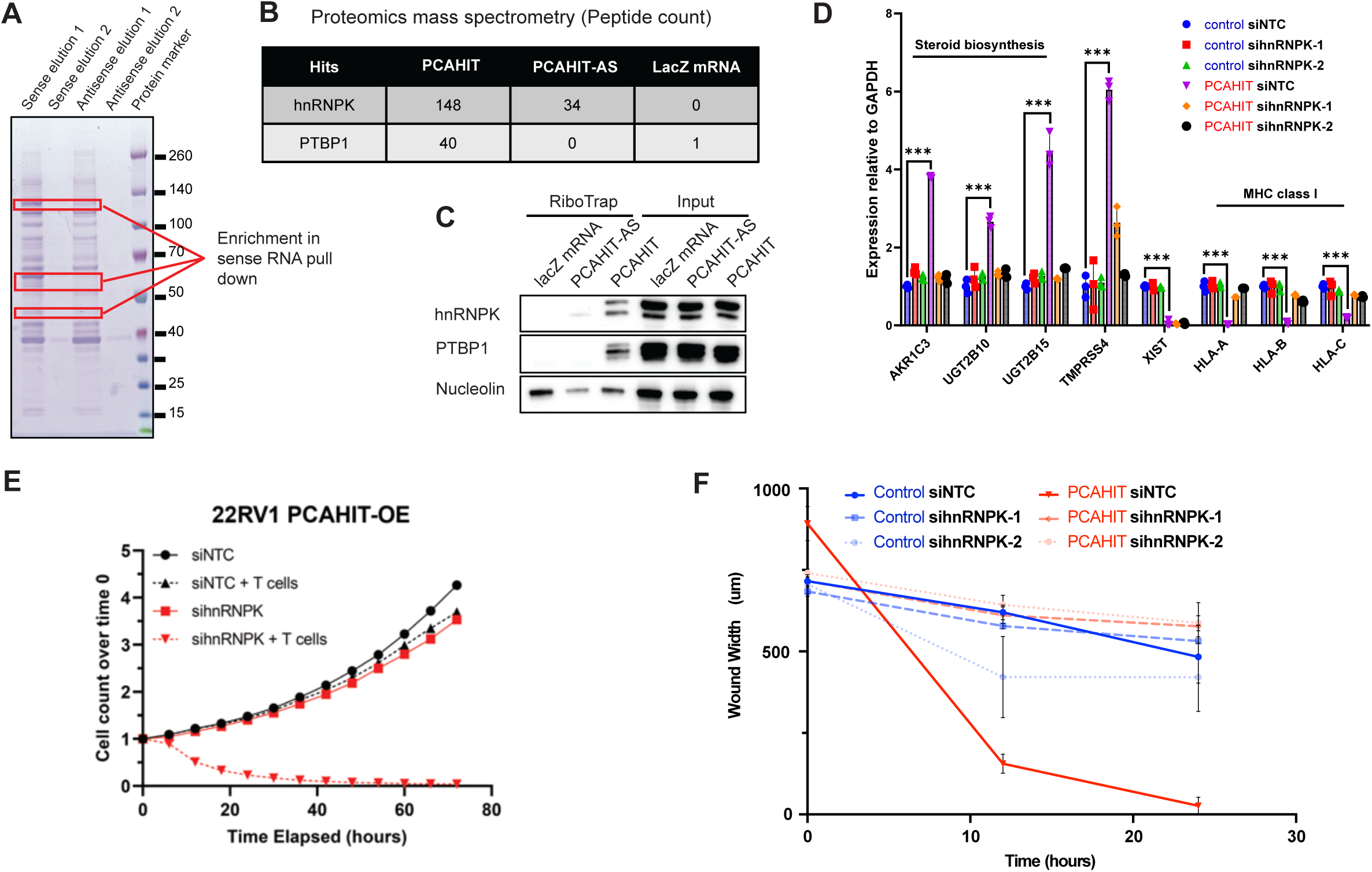
PCAHIT transcriptional activity is mediated by interaction with hnRNPK. (A) Affinity purification using complementary RNA and anti-sense complementary RNA; differentially enriched bands outlined in red. (B) Top two proteins with differential enrichment in sense RNA affinity purification were identified as hnRNPK and PTBP1 using mass spectrometry. (C) RiboTrap affinity assay assessing binding interaction between PCAHIT, PCAHIT anti-sense (AS), and LacZ mRNA with hnRNPK/PTBP1. (D) Expression levels by qPCR of putative PCAHIT regulated genes following PCAHIT overexpression and/or knockdown of hnRNPK (n=3). (E) Proliferation of 22Rv1 cells overexpressing PCAHIT with or without knockdown of hnRNPK and in the presence or absence of T cells (n=3). (F) Quantification of cell migration of 22Rv1 wt or overexpressing PCAHIT with or without knockdown of hnRNPK following mechanical clearance from a portion of a coverslip (n=3).

We then performed knockdown of hnRNPK in the background of PCAHIT overexpression to determine the effect of hnRNPK loss on expression of known PCAHIT target genes. qPCR and western blot analysis confirm efficient loss of mRNA and protein upon siRNA-mediated knockdown (**Supplemental Figure S3D,E**). As described above, PCAHIT overexpression increases expression of steroid hormone biosynthesis genes and decreases expression of HLA class I genes (**Figure 5D**). However, when hnRNPK expression is knocked down, transcription of PCAHIT target genes is returned to baseline (**Figure 5D**), suggesting that hnRNPK is essential for PCAHIT-mediated gene expression regulation. HnRNPK knockdown does not significantly change PCAHIT target gene expression without PCAHIT overexpression, indicating that hnRNPK has no independent regulatory effect under low/absent PCAHIT conditions.

We next sought to determine whether hnRNPK is required not only for the transcriptional outcomes of PCAHIT activity, but for functional outcomes as well. Using a T cell cytotoxicity assay which previously showed decreased T cell-mediated cytotoxicity upon PCAHIT overexpression in 22Rv1 prostate cancer cells, we found that knockdown of hnRNPK restored T cell-mediated cytotoxicity (**Figure 5E**). In addition, using a scratch-wound assay which previously showed increased prostate cancer cell migration upon PCAHIT overexpression, we now find that knockdown of hnRNPK in the context of PCAHIT overexpression reduces cell migration back to baseline (**Figure 5F**). Overall, these findings indicate that knockdown of hnRNPK blocks the transcriptional changes, as well as the phenotype associated with PCAHIT overexpression. This suggests that hnRNPK is a critical mediator of PCAHIT activity, required for PCAHIT-mediated regulation of gene expression.

## Discussion

LncRNAs are increasingly recognized as major drivers of oncogenesis and represent a promising source of novel biomarkers and therapeutic targets. Recent studies have identified several lncRNAs associated with aggressive features and poor clinical outcomes in prostate cancer.(11–15) However, comprehensive annotation of the human genome has identified 58,648 putative lncRNAs,(25) suggesting that the vast majority of lncRNA biology remains unexplored.

In this work, we have identified a novel lncRNA, PCAHIT, which is associated with metastatic recurrence of prostate cancer after prostatectomy. In order to investigate the mechanism by which PCAHIT promotes aggressive prostate cancer phenotypes, we used gene Gene Set Enrichment Analysis (GSEA) on gene expression data from prostate cancer cell lines to show that PCAHIT regulates expression of steroid biosynthesis and MHC Class I genes. We also show that PCAHIT interacts directly with heterogeneous nuclear ribonucleoprotein K (hnRNPK), and that hnRNPK is required for PCAHIT mediated transcriptional modulation. HnRNPK is a member of the hnRNP family of RNA-binding proteins and is known to participate in diverse cellular functions including critical roles mediating the transcriptional regulatory functions of multiple non-coding RNAs.(47)

Consistent with a model in which PCAHIT promotes hnRNPK-dependent transcription of steroid biosynthesis genes leading to resilience in conditions of androgen deprivation as well as downregulation of MHC Class I genes leading to immune evasion, we find that overexpression of PCAHIT causes resistance to T cell-mediated killing and increased proliferation in androgen depleted conditions, which is dependent on expression of hnRNPK. Overall, our data show that PCAHIT drives prostate cancer progression through transcriptional regulation of steroid biosynthesis and immune surveillance pathways and supports further investigation of therapeutic antagonism of PCAHIT and its use as a prognostic biomarker in prostate cancer.

The current work adds to the growing body of evidence suggesting a fundamentally important influence of lncRNA expression on clinical, pathologic, and molecular determinants of prostate cancer aggressiveness. Specifically, prostate cancer is typically dependent on AR signaling for growth and survival, and advanced prostate cancer will often develop resistance to AR-targeted therapies in part through endogenous production of androgens.(36) However, the underlying drivers and mechanisms are poorly understood, which limits opportunities for therapeutic and biomarker development. In addition, most advanced prostate cancers are characterized by an ‘immune cold’ tumor microenvironment,(48) with low levels of infiltrating T cells, low PD-L1 expression, and associated poor response to immune checkpoint inhibitors.(49) Notably, prostate cancer is an outlier among cancer types in that higher levels of infiltrating T cells are associated with worse overall prognosis,(50),(51) implying that T cell activity is inhibited. In particular, MHC-I downregulation is a common way by which multiple cancer types, including prostate cancer, evade T cell-mediated cytotoxicity.(52),(53, 54) However, this process is poorly understood, and to our knowledge, no lncRNA has previously been implicated in regulating MHC-I expression in prostate cancer.

Overall, we find that PCAHIT promotes prostate cancer growth and resistance to AR-targeted therapies through upregulation of endogenous hormone synthesis pathways and downregulation of MHC-I leading to decreased T cell-mediated cytotoxicity. By modulating two distinct pathways promoting prostate cancer growth and treatment resistance, increased PCAHIT expression is a common factor driving these two hallmarks of treatment resistance in advanced prostate cancer. These findings suggest the possibility that PCAHIT may be a useful biomarker of response to anti-androgens in addition to a therapeutic target in prostate cancer using antisense oligonucleotides or small interfering RNAs.

These results also implicate hnRNPK as a key mediator of PCAHIT oncogenic activity. HnRNPK has been previously described as an oncogenic protein and associated with poor outcomes in multiple cancer types, and participates in diverse cellular processes including DNA damage response, glucose metabolism, differentiation, and viral infection.(47) In particular, hnRNPK has a well-described role as a co-regulator of lncRNA activity through multiple general mechanisms including control of mRNA stability and translation, lncRNA localization, lncRNA splicing, genomic organization, and transcriptional output.(47) Further investigation will be required to clarify the specific mechanism by which hnRNPK mediates PCAHIT oncogenic activity.

## Conclusions

In summary, our results reveal a critical role of the novel lncRNA PCAHIT in promoting prostate cancer proliferation, invasion, and metastasis. We further show that PCAHIT promotes transcription of steroid biosynthesis genes and suppresses transcription of MHC-I, resulting in enhanced survival of prostate cancer in androgen depleted conditions, as well as decreased susceptibility to T cell-mediated cytotoxicity, two hallmarks of advanced castration-resistant prostate cancer. Finally, we show that PCAHIT transcriptional regulation depends on the presence of hnRNPK. We expect that these findings will motivate future work to validate PCAHIT as a biomarker of poor outcomes in prostate cancer, and exploration of PCAHIT as a therapeutic target.

## Author Contributions

HL, BM, AMC, JRP, AB, AA, FF, and HL designed research studies; HL, BM, CZ, YN, SZ, KW, SP, SH, TB, PL, MZ, LNC, MC, EAE, JZ, JC, RD, VK, TS, MED, and RM conducted experiments and acquired data; HL, NSY, BM, HJS, CZ, YN, SZ, KW, SP, SH, TB, PL, MZ, LNC, MC, EAE, JZ, JC, RD, VK, TS, MED, and RM analyzed data; TS, MED, and RM provided reagents; NSY, HJS, and HL wrote the manuscript; AA, AMC, FF, and HL supervised the project.

Co-first authors are listed alphabetically.

## Acknowledgments

We would like to acknowledge the following financial support: Prostate Cancer Foundation Young Investigator Award (HL, NSY, SZ, SP, HL, LNC, JC), Department of Defense Prostate Cancer Research Program Early Investigator Research Award (W81XWH-17-1-0192) (HL), Department of Defense Prostate Cancer Research Program Early Investigator Research Award (W81XWH1910424) (SP), Department of Defense Early Investigator award (W81XWH2110046) (LNC), DOD W81XWH2010799 (SZ), NIH 1DP2CA271832-01 (SZ), Benioff Initiative for Prostate Cancer Research (AA), The Parker Institute for Cancer Immunotherapy (AA), National Institutes of Health/National Cancer Institute (K08-CA263552-01A1) (JRP), the V Foundation for Cancer Research [V2024-013] (JRP), JRP is the Ben and Catherine Ivy Foundation Clinical Investigator of the Damon Runyon Cancer Research Foundation [CI-127-24].

## Data Availability Statement

Microarray data is available through the Gene Expression Omnibus (GEO): GSE46691, GSE62116, GSE62667, GSE72291. RNAseq data is available through the Gene Expression Omnibus (GEO): GSE297791. All other data is available upon reasonable request to the corresponding author.

## Ethics Approval Statement

Animal experiments and maintenance were approved by the IACUC of the University of California, San Francisco (protocol AN198293-00Q). Animal studies are reported in compliance with Animal Research: Reporting of In Vivo Experiments (ARRIVE) and the Guide for Care and Use of Laboratory Animals (National Academies Press, 2011).

Human data were previously collected and annotated using a de-identified format. The investigators of the current study had no ability to re-identify participants, and no new data were collected during the current study. Therefore, the current study was determined to not involve human subjects as defined by federal regulations summarized in 45 CFR 46.102(e) and is therefore exempt from University of California, San Francisco Institutional Review Board (IRB) oversight.

## Patient Consent Statement

The requirement for informed consent was waived by the University of California, San Francisco Institutional Review Board (IRB) due to the retrospective nature of the study and the use of de-identified data.

Permission to Reproduce Material from Other Sources: NA; all material is original to the current work.

Clinical Trial Registration: NA; No prospective clinical trial data is included in the current work.

---

Figure S1: Relative expression levels of PCAHIT by qPCR, normalized to GAPDH, in a panel of prostate cancer cell lines (n=3).

Figure S2: qPCR analysis showing expression levels of lncRNAs PCAHIT and SChLAP1 following shRNA-mediated knockdown of PCAHIT using two unique shRNA constructs (n=3).

Figure S3: PCAHIT transcriptional activity is mediated by interaction with hnRNPK. (A) Analysis of microarray gene expression data from six retrospective radical prostatectomy cohorts (Mayo Clinic I, Mayo Clinic II, Cleveland Clinic, Johns Hopkins University, Thomas Jefferson University, and Durham VA) to show metastasis free survival stratified by high/low hnRNPK expression (left) and high/low PTBP1 expression (right). (B) PCAHIT constructs used for affinity purification: Full length (top) and truncated after exon 5 (bottom). (C) PCAHIT affinity purification from LNCaP cells, followed by western blot analysis for hnRNPK. (D-E) Expression levels of hnRNPK detected by western blot (D) and qPCR (E) to confirm siRNA-mediated knockdown of hnRNPK (n=3).

