## Supplementary Material for "Identification and Characterization of PCAHIT, a Novel Long Non-Coding RNA Associated with Poor Outcomes in Prostate Cancer"

#### Figure S1

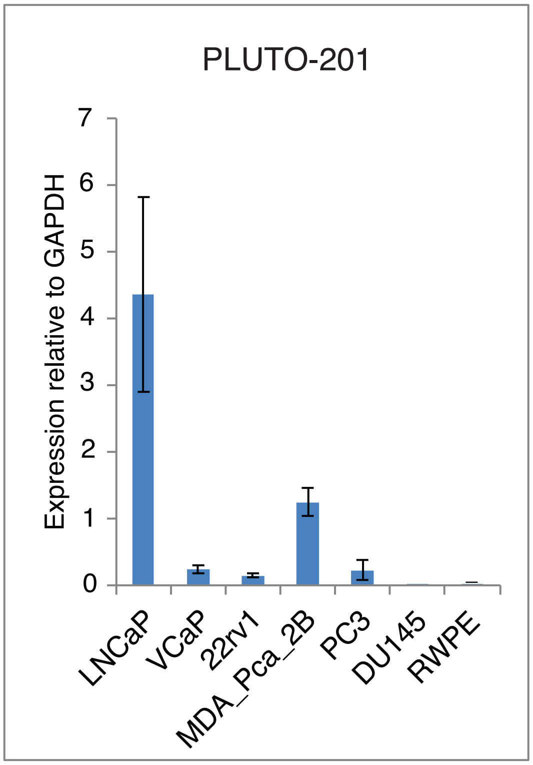


**Figure S1:** Relative expression levels of PLUTO-201 by qPCR, normalized to GAPDH, in a panel of prostate cancer cell lines.

**Figure S2**

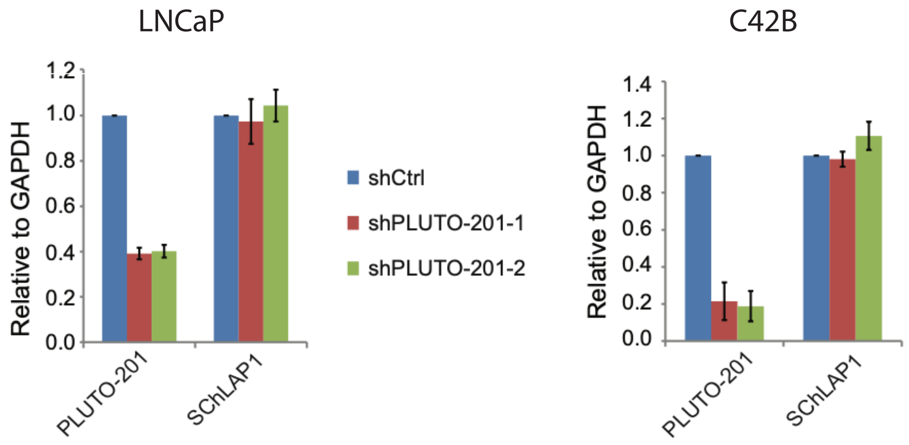


**Figure S2:** qPCR analysis showing expression levels of lncRNAs PLUTO-201 and SChLAP1 following shRNA-mediated knockdown of PLUTO-201 using two unique shRNA constructs.

### Figure S3

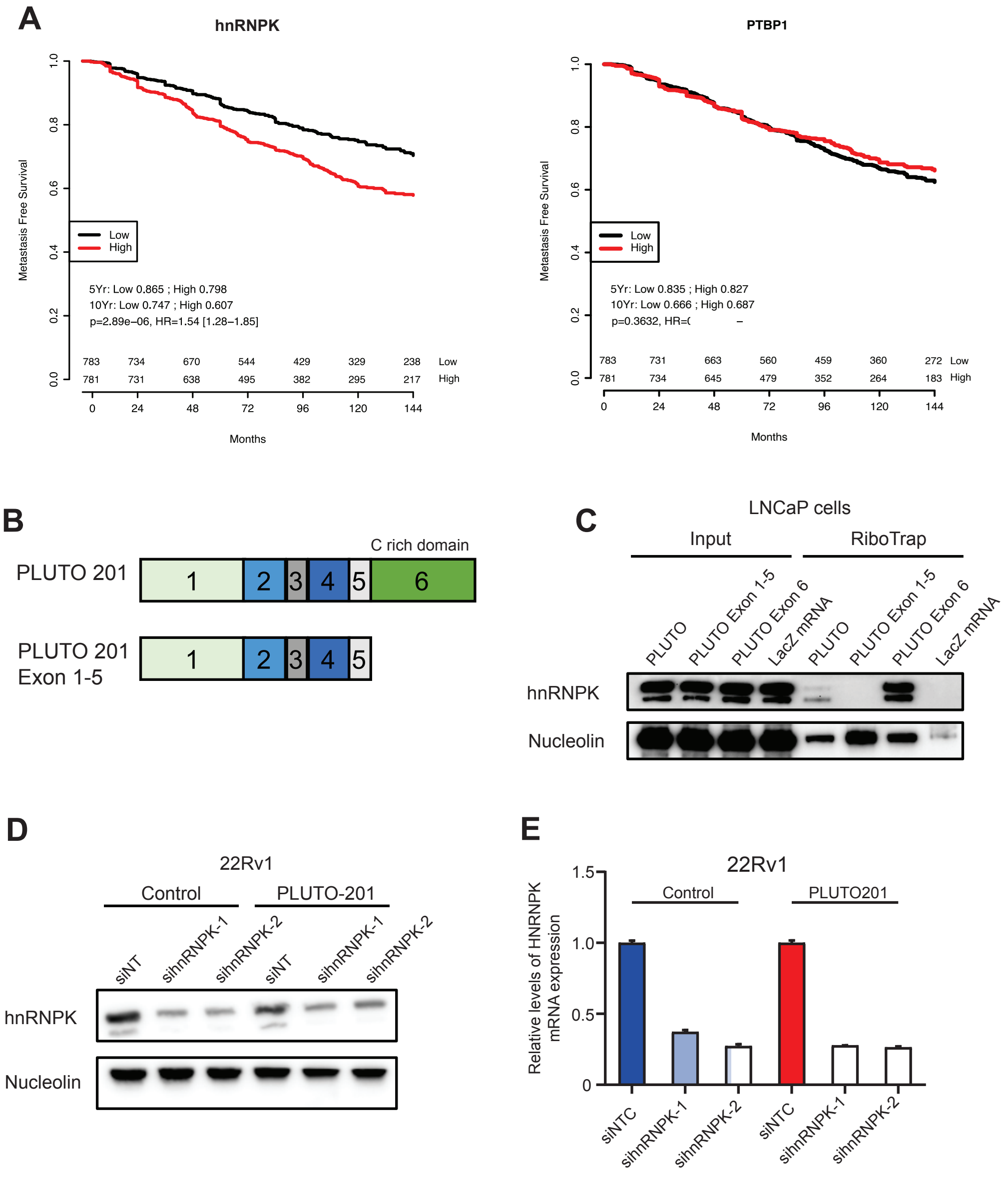


**Figure S3: PLUTO-201 transcriptional activity is mediated by interaction with hnRNPK.** (A) Analysis of microarray gene expression data from six retrospective radical prostatectomy cohorts (Mayo Clinic I, Mayo Clinic II, Cleveland Clinic, Johns Hopkins University, Thomas Jefferson University, and Durham VA) to show metastasis free survival stratified by high/low hnRNPK expression (left) and high/low PTBP1 expression (right). (B) PLUTO-201 constructs used for affinity purification: Full length (top) and truncated after exon 5 (bottom). (C) PLUTO-201 affinity purification from LNCaP cells, followed by western blot analysis for hnRNPK. (D-E) Expression levels of hnRNPK detected by western blot (D) and qPCR (E) to confirm siRNA-mediated knockdown of hnRNPK.

#### Table S1

**smFISH probes targeting PLUTO-201 introns**

ccttgtttctcagagtcaag
aacaagtcagcacacctgag
tgctttagggtcacacaaga
ctgtgtattactcctggatt
gccacattgttgtaacatgt
agcatcagggcattcttaag
gggtaacagtgttatagggg
cacaatcttactctgcctta
tagaagattgcctttggcac
attaactaactgctgcctgt
tgaatccttcccatctttaa
ggcatgaagcaaagatggca
cttccctgaattcaaggtta
tgctccaaagaacattctcc
atctctccctttttttagag
agcttaagtgggggacatta
tgtactattacgtagaccca
tttgcccattaaatatgggc
tatatatctaaacccctcca
agttgcaaaccctgaatgtc
gctatttttaggctcatagc
accttgtgtgaagacgtgac
ttctcacaaatgcaccagtc
aggtattatctacctgggag
aagactggggctgaagcaag
ctgccatatccaaaattgct
agttgtcatgccactaacag
acttaaacacttctctgcct
atatccacagccatcaaact
tacttgatgtccatcttctc
agaagtggcacagtaggttt
aaaatgctgctgctcaggat

**smFISH probes targeting PLUTO-201 exons**

atttcaagtcctcagggtga
cataccacatactgctttca
tgatcagttgaattctctca
gcataaccaaccttctggaa
gtgttgttgttctgaccaat
atggcatgacagaggaaaca
tacattctcttctgcatcag
tcagaagcattttccatact
tttagggctttctgcttgga
attagaaagttttgccatact
tcaggtggatcagttcaagc
